# Serotonin, norepinephrine and acetylcholine differentially affect astrocytic potassium clearance to modulate somatosensory signaling in male mice

**DOI:** 10.1101/2019.12.14.876458

**Authors:** Caitlin A. Wotton, Cassidy D. Cross, Lane K. Bekar

## Abstract

Changes in extracellular potassium ([K^+^]_e_) modulate neuronal networks via changes in membrane potential, voltage-gated channel activity and alteration of transmission at the synapse. Given the limited extracellular space in the CNS, potassium clearance is crucial. As activity-induced potassium transients are rapidly managed by astrocytic Kir4.1 and astrocyte-specific Na^+^/K^+^-ATPase (NKA), any neurotransmitter/neuromodulator that can regulate their function may have indirect influence on network activity. Neuromodulators differentially affect cortical/thalamic networks to align sensory processing with differing behavioral states. Given serotonin (5HT), norepinephrine (NE), and acetylcholine (ACh) differentially affect spike frequency adaptation and signal fidelity (“signal-to-noise”) in somatosensory cortex, we hypothesize that [K^+^]_e_ may be differentially regulated by the different neuromodulators to exert their individual effects on network function. This study aimed to compare effects of individually applied 5HT, NE, and ACh on regulating [K^+^]_e_ in connection to effects on cortical evoked response amplitude and adaptation in male mice. Using extracellular field and K^+^ ion-selective recordings of somatosensory stimulation, we found that differential effects of 5HT, NE, and ACh on [K^+^]_e_ regulation mirrored differential effects on amplitude and adaptation. 5HT effects on transient K^+^ recovery, adaptation and field post-synaptic potential amplitude were disrupted by barium (200 µM), whereas NE and ACh effects were disrupted by ouabain (1 µM) or iodoacetate (100 µM). Considering the impact [K^+^]_e_ can have on many network functions; it seems highly efficient that neuromodulators regulate [K^+^]_e_ to exert their many effects. This study provides functional significance for astrocyte-mediated buffering of [K^+^]_e_ in neuromodulator-mediated shaping of cortical network activity.

**Significance statement:** We demonstrate that the neuromodulators serotonin, norepinephrine, and acetylcholine all have distinct effects on astrocyte-mediated extracellular potassium regulation and that these differential actions are associated with the different effects of the neuromodulators on cortical networks. By affecting astrocytic potassium regulation, long-range neuromodulatory networks can rapidly and efficiently affect broad areas of the brain. Given that neuromodulatory networks are at the core of our behavioral state and determine how we interact with our environment, these studies highlight the importance of basic astrocyte function in general cognition and psychiatric disorders.

## Introduction

Changes in extracellular potassium concentration ([K^+^]_e_) modulate neuronal networks via membrane depolarization/hyperpolarization with subsequent activation/inactivation of voltage-gated channels and alteration of transmission at the synapse(Bellot-Saez, Kekesi, Morley, & Buskila, 2017; Kofuji & Newman, 2004; Larsen, Stoica, & MacAulay, 2016; Sibille, Dao Duc, Holcman, & Rouach, 2015; Sibille, Pannasch, & Rouach, 2014). Relatively limited extracellular space in the CNS increases potential for small changes in [K^+^]_e_ to have a significant impact on neuronal activity. Therefore, it is crucial to clear potassium rapidly and efficiently. It is now widely accepted that activity-induced changes in [K^+^]_e_ are rapidly managed by the astrocytic potassium inward rectifying channel subtype 4.1 (Kir4.1)(Bellot-Saez et al., 2017; Butt & Kalsi, 2006; Chever, Djukic, McCarthy, & Amzica, 2010; D’Ambrosio, Gordon, & Winn, 2002; Larsen et al., 2014; Larsen & MacAulay, 2014; Sibille et al., 2015) and the astrocyte-specific α2β2 isoform of the Na^+^/K^+^-ATPase (NKA)(D’Ambrosio et al., 2002; Larsen et al., 2014; Pellerin & Magistretti, 1997; Stoica et al., 2017). Due to the highly negative resting membrane potential of astrocytes, Kir4.1 is highly responsive to increases in [K^+^]_e_ with local inward K^+^ uptake into astrocytes(Sibille et al., 2015; Sibille et al., 2014) whereas the astrocyte-specific NKA is additionally responsive to changes in intracellular Na^+^ driven by the synapse-dependent glutamate transporter(Larsen, Holm, Vilsen, & MacAulay, 2016; Larsen, Stoica, et al., 2016; Pellerin & Magistretti, 1997) and/or the Na^+^/Ca^2+^ exchanger in response to astrocyte Ca^2+^ transients(Wang, Smith, et al., 2012). Accordingly, any neurotransmitter or neuromodulator that can alter or regulate astrocytic Kir4.1 or NKA activity may have indirect influence on network activity/function via [K^+^]_e_.

Neuromodulators are known to be involved in regulating cortical activity via both local effects at the cortical level as well as changes at the level of the thalamus(McCormick, 1992). Differential neuromodulator effects on cortical and thalamic networks may align sensory processing with different behavioral states(Castro-Alamancos & Gulati, 2014). Norepinephrine (NE), acetylcholine (ACh) and serotonin (5HT) all appear to decrease spike frequency adaptation (referred to as ‘adaptation’ from here out) in part via blockade of various potassium conductances (*I*_M_ and *I*_AHP_)(McCormick, 1992), whereas effects on signal fidelity (“signal-to-noise”) in cortical layer II/III results from differing effects on corticothalamic/thalamocortical interactions(Castro-Alamancos & Gulati, 2014) and/or effects on local inhibitory circuits (Deng & Lei, 2008; Lei, Deng, Porter, & Shin, 2007; McCormick, 1992; Salgado et al., 2011; Xiao, Deng, Yang, & Lei, 2009). Local inhibitory networks are most relevant in isolated slice models and are sensitive to both purinergic and astrocyte-directed pharmacology(Quon, Wotton, & Bekar, 2017; Wotton, Quon, Palmer, & Bekar, 2018). A recent study by Ding et al. (2016) demonstrated that a cocktail of neuromodulators containing NE, ACh, dopamine, orexin, and histamine induce a wakefulness brain state via an increase in [K^+^]_e_(Ding et al., 2016; Rasmussen et al., 2019). We suggest that, given the differential effects of neuromodulators on circuit function, [K^+^]_e_ may be differentially regulated by the different neuromodulators. This study aimed to compare and contrast the effects of NE, ACh, and 5HT on regulating [K^+^]_e_ in connection to effects on cortical evoked response amplitude and adaptation. Using extracellular field and K^+^ ion-selective microelectrode recordings in response to stimulation trains in somatosensory layer IV-II synapses, we found that the differential effects of NE, ACh, and 5HT on [K^+^]_e_ regulation mirrored the differential effects on amplitude and adaptation. Furthermore, we show that 5HT-mediated effects are dependent on Ba^2+^-sensitive Kir channels and NE- and ACh-mediated effects are dependent on opposing action on the ouabain-sensitive NKA.

## Materials and Methods

### Slice preparation

A total of 105 male C57Bl6 mice aged 40-70 days were used in these studies. Although no sex differences were anticipated, males were used in these first studies to avoid potential variation due to staging differences of the estrous cycle in females. Mice were housed two per cage and kept on a 12:12 light:dark schedule until used for experiments. For acute slice preparation, mice were anesthetized with isoflurane and sacrificed by decapitation. Brains were dissected out and placed in ice cold slicing solution consisting of (in mM): 130 Choline Chloride, 3 KCl, 1.25 NaHPO_4_, 2 CaCl_2_, 2 MgSO_4_, 24 NaHCO_3_, 10 dextrose, and 1 lactate. The brains were sliced coronally (350 µM) with a vibratome (Leica 1200) in the same ice-cold solution and subsequently placed in a chamber containing normal artificial cerebral spinal fluid (aCSF) bubbling with 95% oxygen 5% CO_2_ and kept at 32°C for 1 hour. Normal aCSF is the same as cutting solution but with 130 NaCl replacing 130 Choline Chloride. Following the hour at 32°C, the slices were then kept at room temperature for another hour before beginning experiments. All experiments were conducted at room temperature unless noted and in accordance with the guidelines outlined by the Canadian Council on Animal Care and were approved by the University of Saskatchewan Committee on Animal Care and Supply.

### Ion-selective electrodes and Extracellular field recordings

Electrodes were prepared daily by the following procedure: 1) borosilicate glass capillaries (WPI) pulled on a Narishige PC-10 puller (∼5 megaohms) are filled using negative-pressure with HCl for one minute to expose the glass molecules for strong silane binding, 2) electrode is rinsed with acetone and the inner layer of the pipette tip is coated with silane (Sigma) and placed on hot plate (pre-heated to 280°C) for 10 minutes. 3) The electrode is then backfilled with 150 mM KCl and negative-pressure used again to pull ionophore complex B (Sigma) into the tip (300-400 µm length). This valinomycin ionophore is the most selective K^+^ exchanger available (Ammann, Chao, & Simon, 1987). A calibration was performed prior to and following each experiment in the bath perfusate using aCSF containing 2, 3, and 6 mM potassium. Any ion-selective electrodes showing less than 5 mV voltage change in response to 1 mM change in potassium were excluded.

Slices were placed in a perfusion chamber on a Zeiss Axioskop upright microscope that was perfused at a rate of ∼4 mL per minute with normal aCSF bubbled with 95% O_2_ and 5% CO_2_. Slices were maintained in position using a slice holder (Warner Instruments). Experiments were performed by first placing the reference/field electrode (filled with 0.9% saline) in layer II of hindlimb/forelimb somatosensory cortex for recording using a differential amplifier (DP311; Warner Instruments) attached to a digidata 1440A (Molecular Devices). Signals were captured at a rate of 2 KHz using a highpass filter of 0.1 Hz and lowpass filter of 1000 Hz for field recordings or low pass digital filter of 10 Hz for ion-selective reference. The stimulating electrode (tungsten concentric bipolar, TM33CCINS, WPI) was placed in layer IV and driven by an Iso-Flex stimulator (AMPI, Isreal). After achieving an acceptable extracellular field recording (maximum amplitude of at least 1.2 mV), the ion-selective electrode attached to an Axoprobe-1A multielectrode amplifier (high resistance head-stage and 10 Hz lowpass filter) was then placed within 10 µm of the reference/field electrode. A ten-pulse stimulation protocol was repeated every 2 minutes for assessment of evoked potassium responses. For quantification of evoked response recovery tau, individual responses were fit using the exponential, sloping baseline function in Clampfit 10 software (Chebyshev method).

### Drugs

Acetylcholine (50 µM; Sigma), BaCl_2_ (200 µM; Sigma), iodoacetate (100 µM; Sigma), norepinephrine hydrochloride (50 µM; Sigma), ouabain tetrahydrochloride (1 µM; Sigma), serotonin hydrochloride (20 µM; Sigma), tetrodotoxin (200 ηM; HelloBio).

### Statistics

All data are expressed as mean ± SD and were compared using repeated measures two-way or three-way analysis of variance (Prism v 8.1.2, GraphPad Software, CA) to assess neuromodulator and pharmacology treatment effects and interactions (sphericity was assumed). All experimental sets were paired, enabling normalization of effects to percentage of baseline values in some instances. At least 3 stimuli/protocols (amplitude or recovery tau) were averaged before and after drug administration for comparison. Fisher’s least significant difference post-hoc was used to compare differences between groups. Number of animals used in each set of experiments are indicated in parentheses in all figures. All experimental sets were obtained from a minimum of 3 separate animals with a maximum of 3 slice experiments from a given animal.

## Results

### Neuromodulators differentially affect regulation of extracellular potassium

As a previous study showed that a cocktail of neuromodulators could alter extracellular potassium to affect brain state(Ding et al., 2016), we chose to explore and characterize serotonin, norepinephrine and acetylcholine effects on cortical extracellular potassium responses independently. To achieve this, we placed an ion-sensitive microelectrode within 10 µm of the field recording electrode (which also served as a reference for the potassium recordings) in cortical layer II (Fig. 1) and followed the height of the evoked K^+^ response that occurred following the 10-pulse (20 Hz) stimulation applied every two minutes (10-pulse stimulation embedded in rising phase of evoked response, Fig. 1), the decay time constant (tau) of the stimulated K^+^ response recovery, and the baseline [K^+^]_e_. The recovery tau of the evoked response was decreased (indicating a faster recovery back to basal K^+^ level) by both 5HT and NE, but not affected by ACh (Fig. 2A; NM x time interaction p = 0.040; F_2, 33_ = 3.554). The amplitude of the evoked K^+^ response decreased in response to each neuromodulator (Fig. 2B; NM x time interaction p = 0.020; F_2, 32_ = 4.403). Given that ACh demonstrated a decrease in the evoked K^+^ response similar to that of 5HT and NE but did not alter the recovery tau, it is unlikely that the change in recovery tau is dependent on the amplitude of the small evoked responses (0.3-0.8 mM) under the conditions used in these studies. Consistent with this, no correlation was found between change in recovery tau and change in evoked response amplitude (pearson r = −0.215, R^2^ = 0.0462, P = 0.189, n = 39). The basal concentration of K^+^ was increased with application of 5HT and NE while ACh had no effect (Fig. 2C, D top; NM x time interaction p = 0.020; F_2, 23_ = 4.678). However, with the understanding that these neuromodulators each affect excitability and thus spontaneous activity in the slice that could impact baseline [K^+^]_e_, we also assessed the neuromodulator effects on baseline [K^+^] in the presence of 200 ηM tetrodotoxin (TTX; Na channel blocker) to inhibit all neural activity. Note the loss of evoked responses during wash-in of TTX prior to application of the different neuromodulators (Fig. 2D bottom). In the absence of effects on neural activity, the neuromodulators demonstrate significantly different effects on baseline [K^+^]. 5HT shows no influence, NE shows a decrease and ACh shows an increase (Fig. 2D bottom, E; NM x time interaction p < 0.001; F_2, 14_ = 22.15). Thus, only 5HT and NE affect an increase in recovery of evoked K^+^ responses while NE also decreases and ACh only increases baseline [K^+^].

**Figure 1:**
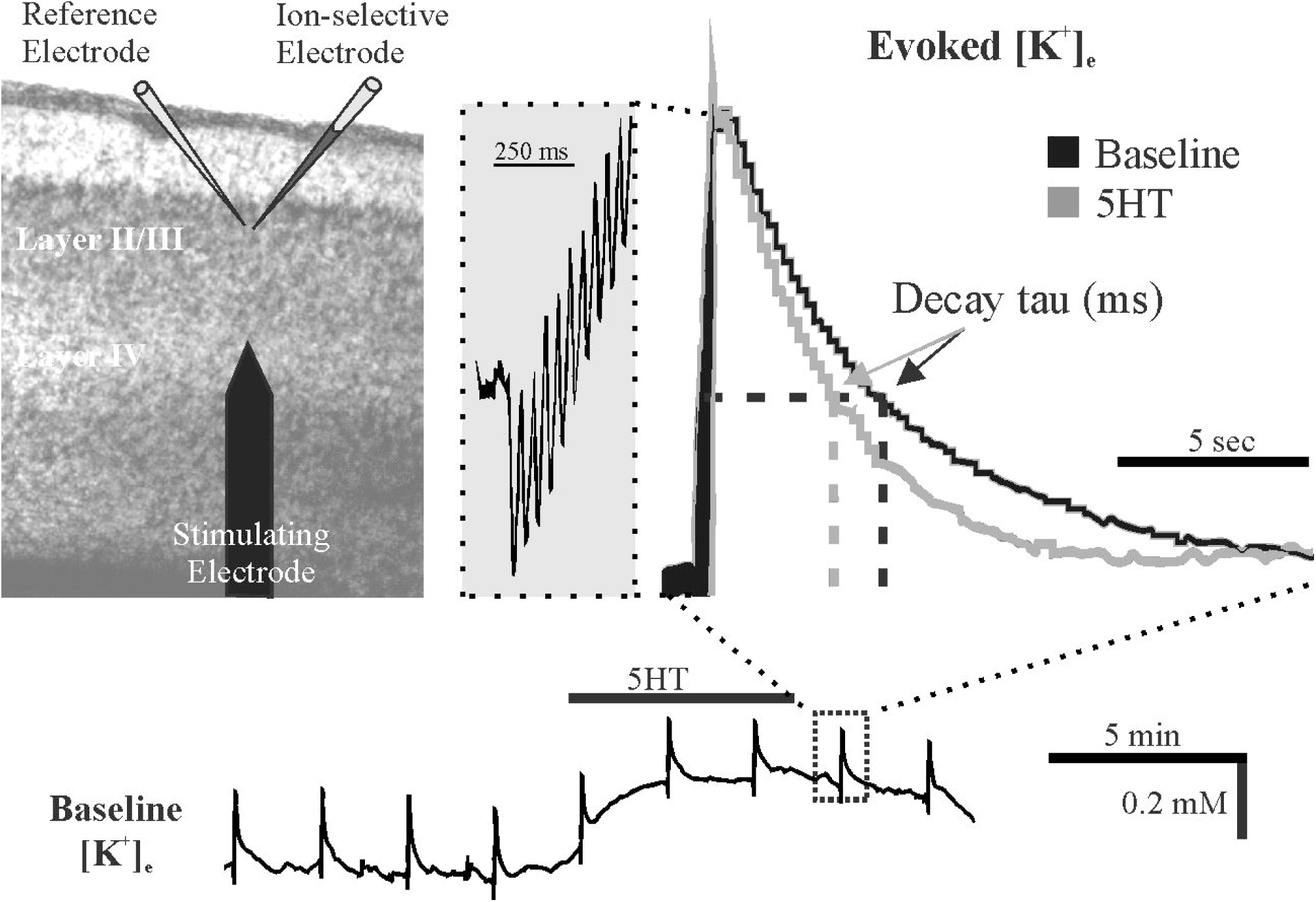
Experimental design. The stimulating electrode is placed in layer IV of somatosensory cortex and the reference/K^+^ ion-sensitive electrodes are placed within 10 µm of each other in layer II. At the beginning of each 2-minute sweep, 10 pulses (20 Hz) stimulate a small evoked K^+^ response. Extracellular K^+^ level is continuously monitored for the duration of recording. Administration of 5HT is shown as an example where 5HT increases the baseline K^+^ (bottom trace) and decreases the recovery tau (enlarged overlayed traces).

**Figure 2:**
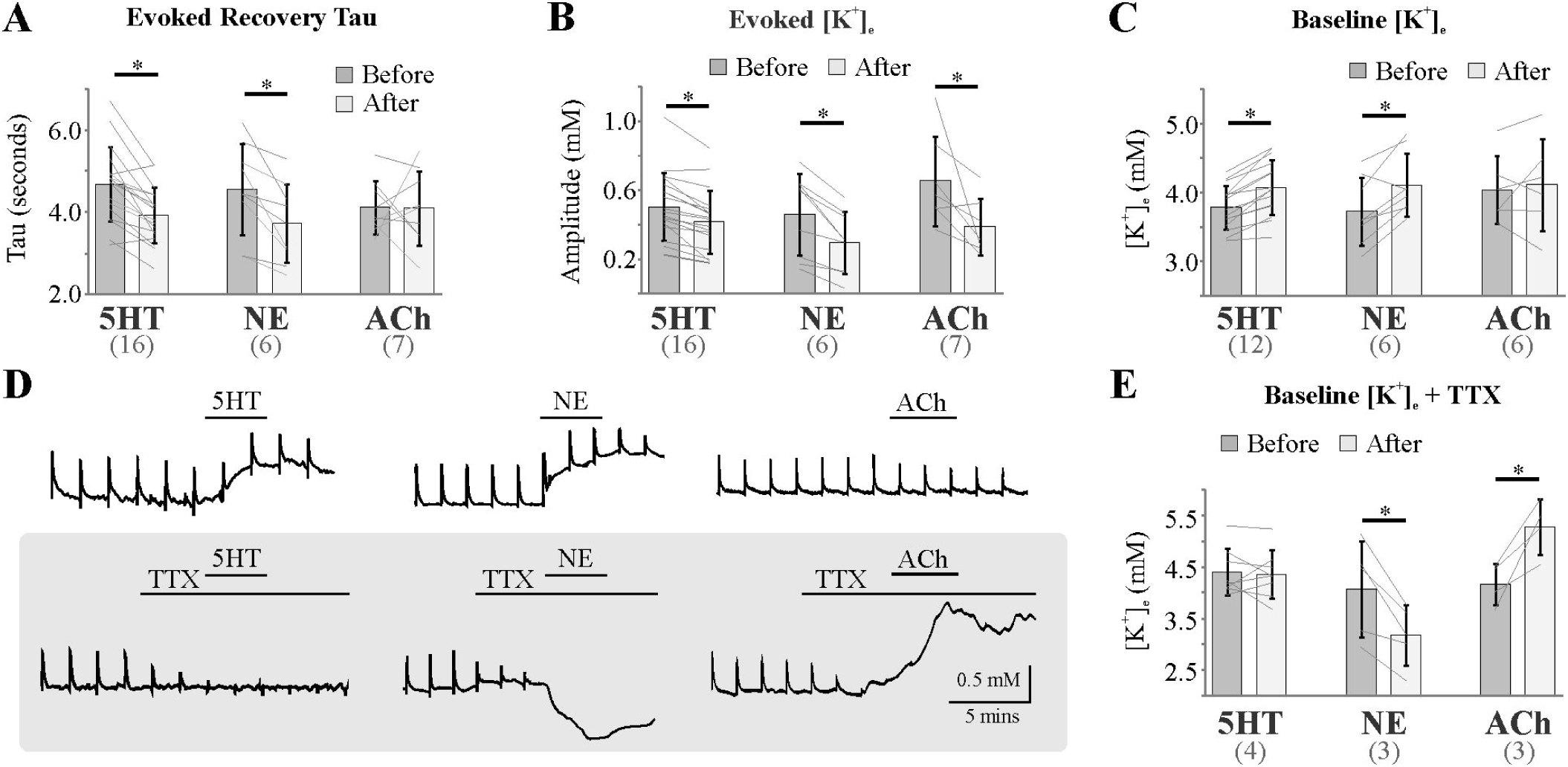
Neuromodulators differentially affect K^+^ homeostasis. **A)** Recovery tau was decreased in response to 5HT and NE, but not altered by ACh. **B)** The evoked transient K^+^ response was decreased by all neuromodulators. **C)** 5HT and NE displayed an increase in the basal K^+^ level whereas ACh did not. **D** and **E)** Neuromodulator responses on basal K^+^ levels differ substantially when neural activity is blocked with TTX. Representative traces showing neuromodulator effects on basal K^+^ without (top) or with (bottom) 200 ηM TTX. * p < 0.05, two-way RM ANOVA comparisons with Fisher’s LSD. n = 4-19 slice experiments. Numbers in parentheses on x-axis indicate the number of animals.

### Neuromodulators affect K^+^ regulation via Kir4.1 and the Na^+^/K^+^-ATPase

The inwardly rectifying K^+^ channel (Kir4.1) and Na^+^/K^+^-ATPase (NKA) are proposed as the major means through which extracellular K^+^ is cleared from the synapse by astrocytes (Chever et al., 2010; Larsen et al., 2014; Larsen, Stoica, et al., 2016; Sibille et al., 2015; Sibille et al., 2014). To assess the role of Kir4.1 and NKA in recovery from transient increases as well as determining basal K^+^ levels, we used barium chloride (BaCl_2_; Ba^2+^; 200 µM) and ouabain (1 µM) to inhibit the Kir4.1 and NKA, respectively. This low dose of ouabain is effective in blocking the α2 and α3 isozymes of the NKA (Sweadner, 1989), the α2 being selective to astrocytes with kinetics ideal for responding to transient changes in extracellular K^+^ (Larsen et al., 2014). We observed a total block of the 5HT-mediated decrease in the evoked recovery tau with both Ba^2+^ and ouabain pre-treatments (Fig. 3A, left; pharm x time interaction p = 0.010; F_2, 29_ = 5.475). Although the interaction did not reach significance (pharm x time interaction p = 0.088; F_2, 15_ = 2.863), NE showed a similar loss of effect from baseline on evoked recovery tau when in the presence of Ba^2+^ or ouabain (Fig. 3A, right).

**Figure 3:**
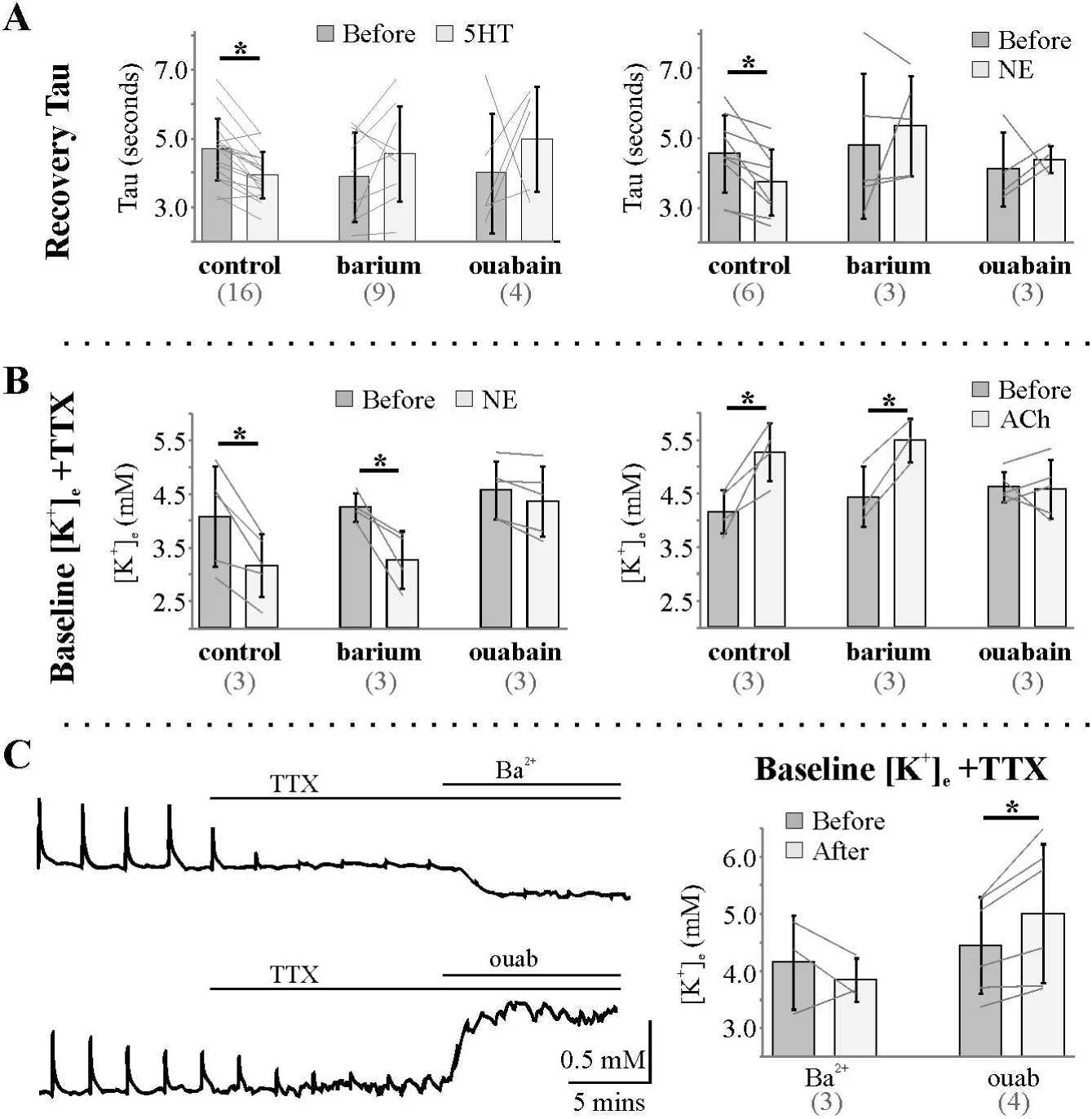
Neuromodulator effects on extracellular K^+^ are susceptible to blockade of Kir4.1 channels and the sodium-potassium ATPase. **A)** The 5HT- and NE-mediated decrease in the recovery tau are lost in the presence of either antagonist. As ACh did not impact recovery tau, it was not assessed pharmacologically. **B)** The NE- and ACh-mediated effects on baseline K^+^ in the presence of TTX were not affected by Ba^2+^ but completely blocked in low-dose ouabain. **C)** Ba^2+^ did not affect baseline potassium whereas ouabain showed an increase. * p < 0.05 two-way RM ANOVA with Fisher’s LSD. n = 3-18. Numbers in parentheses on x-axis indicate the number of animals.

The impact of the antagonists on basal [K^+^] were studied after block of synaptic/neural activity with the Na^+^ channel blocker TTX (200 ηM, Fig. 3B). Interestingly, while Ba^2+^ had no effect, low-dose ouabain effectively blocked both the NE-mediated decrease (pharm x time interaction p = 0.031; F_2, 11_ = 4.871) and the ACh-mediated increase (pharm x time interaction p = 0.009; F_2, 9_ = 8.171) in basal [K^+^]_e_, suggesting opposing effects on the NKA (Fig. 3B). As ouabain wash-in increases basal [K^+^]_e_ similarly to ACh (Fig. 3C), it may be that ouabain occludes an ACh-mediated inhibition of NKA activity, whereas it inhibits an NE-mediated activation. Barium notably, does not impact basal [K^+^]_e_ levels on its own (Fig. 3C).

### Neuromodulators differentially affect synaptic adaptation and amplitude

In attempt to establish a physiological relevance for neuromodulator-mediated effects on extracellular [K^+^], we assessed the effects of the neuromodulators on adaptation and amplitude of field excitatory post-synaptic potentials (fEPSPs) in our somatosensory cortical model. In the cortex in response to trains of incoming information, the amplitude of the first fEPSP is largest with subsequent fEPSPs rapidly decreasing through a process termed adaptation. At high frequency it is thought that neurotransmitter depletion plays a dominant role whereas at lower frequencies many factors appear to be involved (Abbott, Varela, Sen, & Nelson, 1997; Fioravante & Regehr, 2011; Varela et al., 1997; Zucker & Regehr, 2002). In addition to effects on the degree of somatosensory adaptation, neuromodulators are also known to impact the signal-to-noise ratio of somatosensory circuits (Castro-Alamancos & Gulati, 2014; Hirata, Aguilar, & Castro-Alamancos, 2006; McCormick, 1992) that can be observed in our cortical slice model (simplistically) as a change in overall amplitude of the fEPSPs. Field recordings do not enable evaluation of the full complexity of changes in the circuit as they do not differentiate effects on excitatory vs inhibitory activity. However, the output can still be appreciated in general amplitude changes.

We assess neuromodulator-mediated effects on adaptation using a low-frequency stimulation protocol (10 pulses at 20 Hz) in extracellular field recordings (Fig. 4A). All trains of 10 pulses had fEPSP amplitudes normalized to the first fEPSP in the train so adaptation responses could be displayed (Fig. 4B) and directly compared across groups (Fig. 4C). For ease of comparison, a single adaptation value (fraction of first fEPSP amplitude) was determined for each train as the average of normalized pulses 2-10 (Fig 4C insets). Both 5HT and NE decrease adaptation (increase amplitude) of fEPSPs 2-10 or 3-10, respectively (Fig. 4C). In contrast, ACh has no impact on the degree of adaptation in our somatosensory slice model. Thus, the neuromodulators differentially affect adaptation (NM x time interaction p = 0.002; F_2, 30_ = 7.764). Effects of neuromodulators on the first fEPSP amplitude are presented as fold change of baseline recordings immediately prior to neuromodulator application (Fig. 4D). All three neuromodulators decrease the amplitude of fEPSPs (neuromodulator effect P < 0.001; F_1, 30_ = 34.95). Notably, the differential effect on adaptation mirrors the differential effect observed on evoked K^+^ recovery tau (Fig. 2A), raising the question of whether [K^+^]_e_ regulation is involved in adaptation responses.

**Figure 4:**
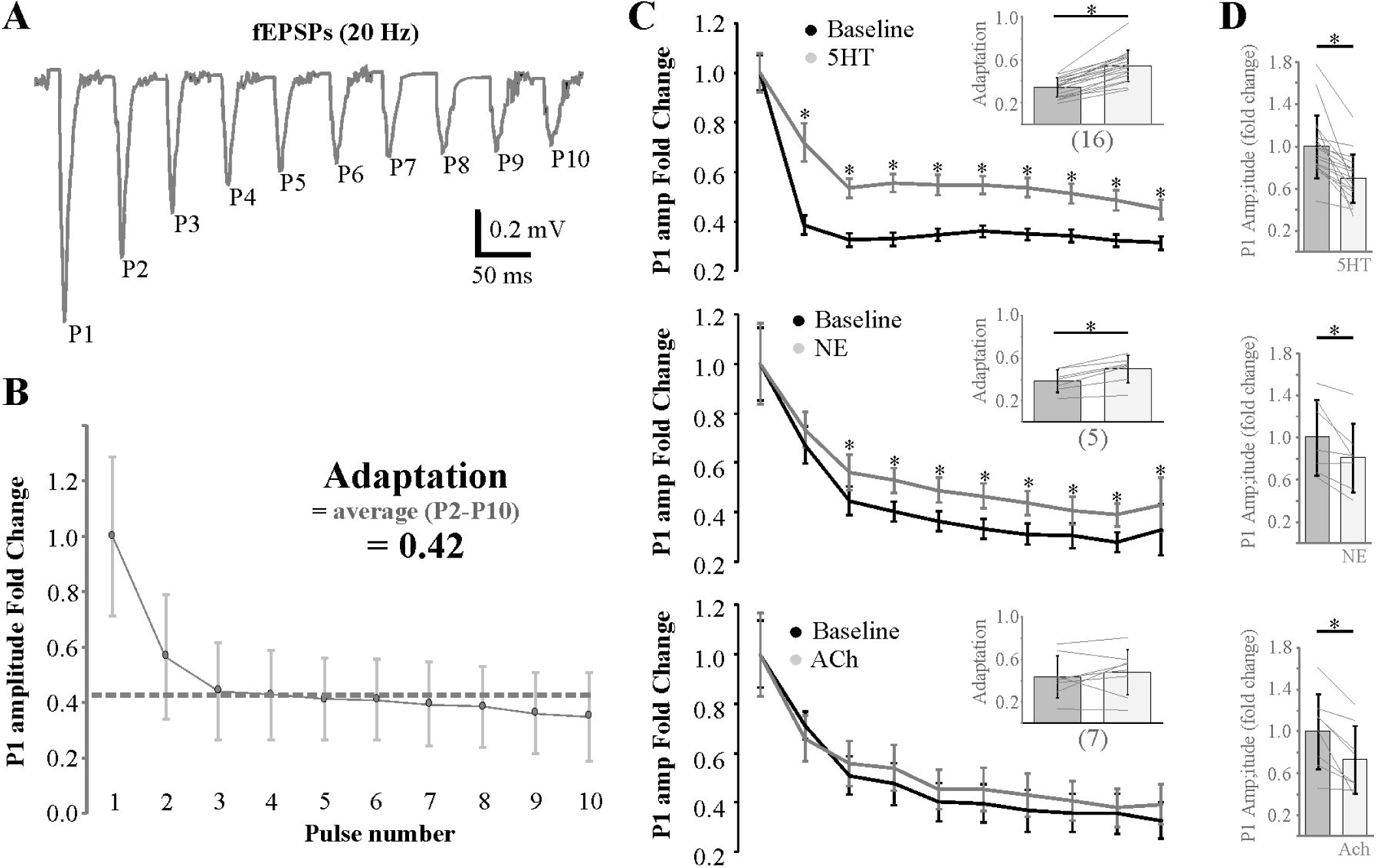
Neuromodulators differentially affect cortical somatosensory adaptation. **A)** Adaptation was assessed from extracellular field recordings of layer IV-II synapses using a 10-pulse protocol at 20 Hz. **B)** Pulse amplitude was normalized to the first pulse and adaptation is quantitated as the average of pulses 2-10 for each train. **C)** Note that whereas 5HT and NE both decrease adaptation, ACh as no effect. D) Despite affecting adaptation differently, all three neuromodulators decrease the amplitude of P1 upon administration. * p < 0.05 two-way RM ANOVA with Fisher’s LSD. n = 6-18. Numbers in parentheses on x-axis indicate the number of animals.

### Dysregulation of K^+^ impedes neuromodulator effects on adaptation and amplitude

If changes in regulation of [K^+^]_e_ are involved in neuromodulator-mediated somatosensory adaptation, it would stand to reason that pharmacological disruption of neuromodulator effects on K^+^ homeostasis should also disrupt adaptation. To assess this possibility, 5HT and NE effects on adaptation were repeated in the presence of Ba^2+^ or ouabain (Fig. 5), both of which affected neuromodulator-mediated effects on K^+^ homeostasis (Fig. 3). 5HT effects on adaptation were inhibited in the presence of Ba^2+^, but unaffected in the presence of ouabain (Fig. 5, left; pharm x time interaction p < 0.001; F_2, 32_ = 12.03). In contrast, NE effects on adaptation were inhibited in the presence of ouabain, but unaffected in the presence of Ba^2+^ (Fig. 5, right; pharm x time interaction p = 0.427; F_2, 14_ = 0.906). Although post-hoc comparisons did not demonstrate differences before and after NE in the presence of Ba^2+^ (Fig. 5, top right), two-way repeated measures ANOVA supported a significant NE effect (p = 0.017; F_1, 50_ = 6.091); consistent with the typically smaller effect of NE on adaptation than 5HT under control conditions (Fig. 4C). The differential effects of K^+^-directed pharmacology on 5HT and NE are confirmed by a three-way repeated measures ANOVA (pharm x NM x time interaction p = 0.001; F_2, 44_ = 7.833). This data indicates that inward-rectifiers play a role in the 5HT-mediated decrease in adaptation, while the NE-mediated decrease was dependent on NKA activity.

**Figure 5:**
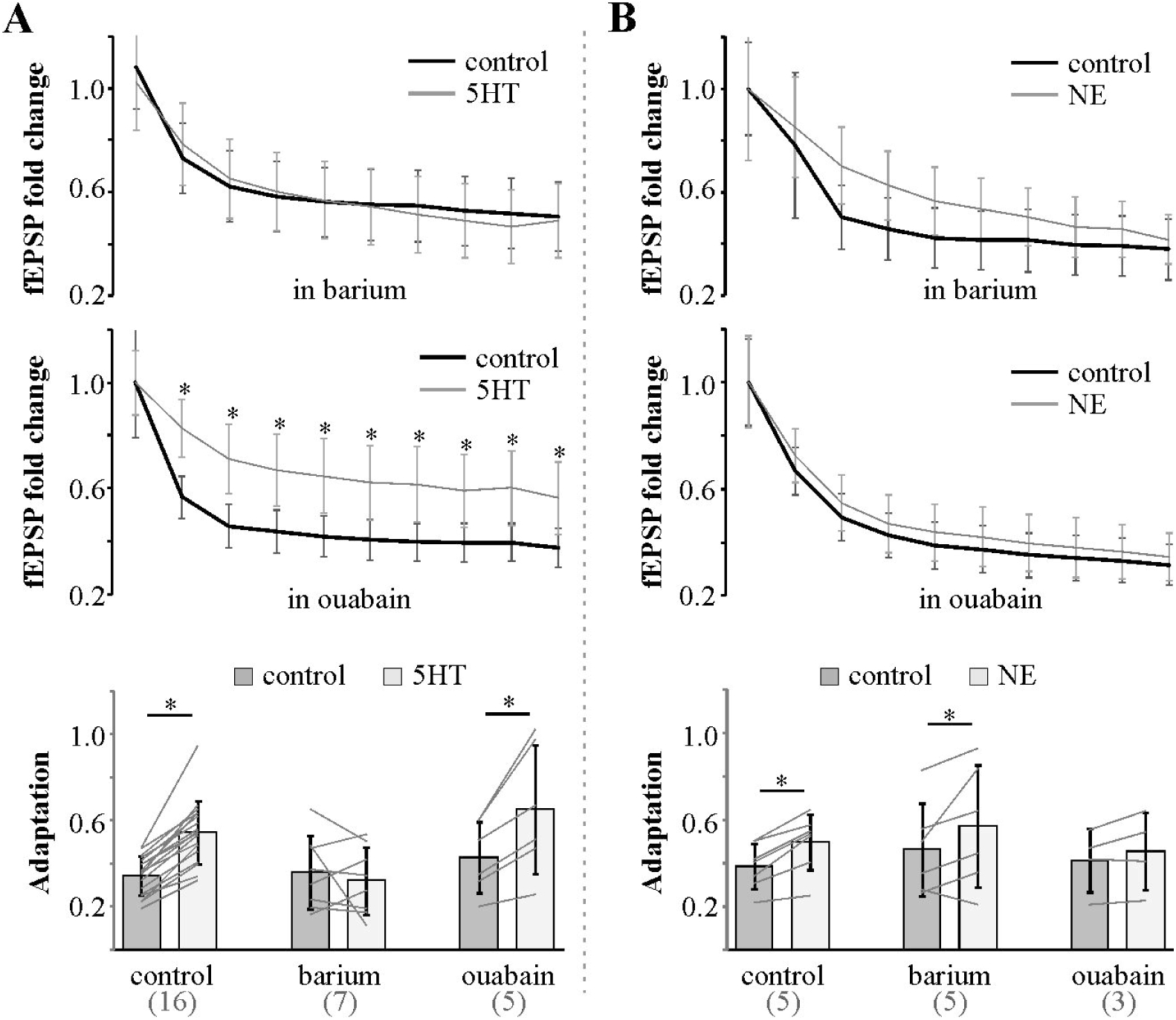
Pharmacological disruption of K^+^ regulation differentially affects neuromodulator-mediated adaptation. **A)** 5HT-mediated changes in adaptation were abolished with Ba^2+^ but not low-dose ouabain. **B)** In contrast, NE-mediated changes to adaptation were abolished by low-dose ouabain, but not Ba^2+^. * p < 0.05 two-way repeated measures ANOVA with Fisher’s LSD. n = 4-18. Numbers in parentheses on x-axis indicate the number of animals.

Whereas the 5HT-mediated decrease of fEPSP amplitude is inhibited by Ba^2+^ and not ouabain in the perfusate, the NE- and ACh-mediated decreases are inhibited by ouabain and not Ba^2+^ in the perfusate (Fig. 6A-C). These results align well with the 5HT-mediated effects on [K^+^] regulation being primarily dependent on the Ba^2+^-sensitive Kir4.1 and both NE and ACh-mediated effects dependent on the ouabain-sensitive NKA (Fig 3). Likewise, as ouabain affected fEPSP amplitude upon washin (Fig. 6D; associated with increase in [K^+^]_e_, Fig. 3C), it is possible that it served to occlude ACh-mediated inhibition of NKA activity and inhibit NE-mediated activation to impact effects on amplitude.

**Figure 6:**
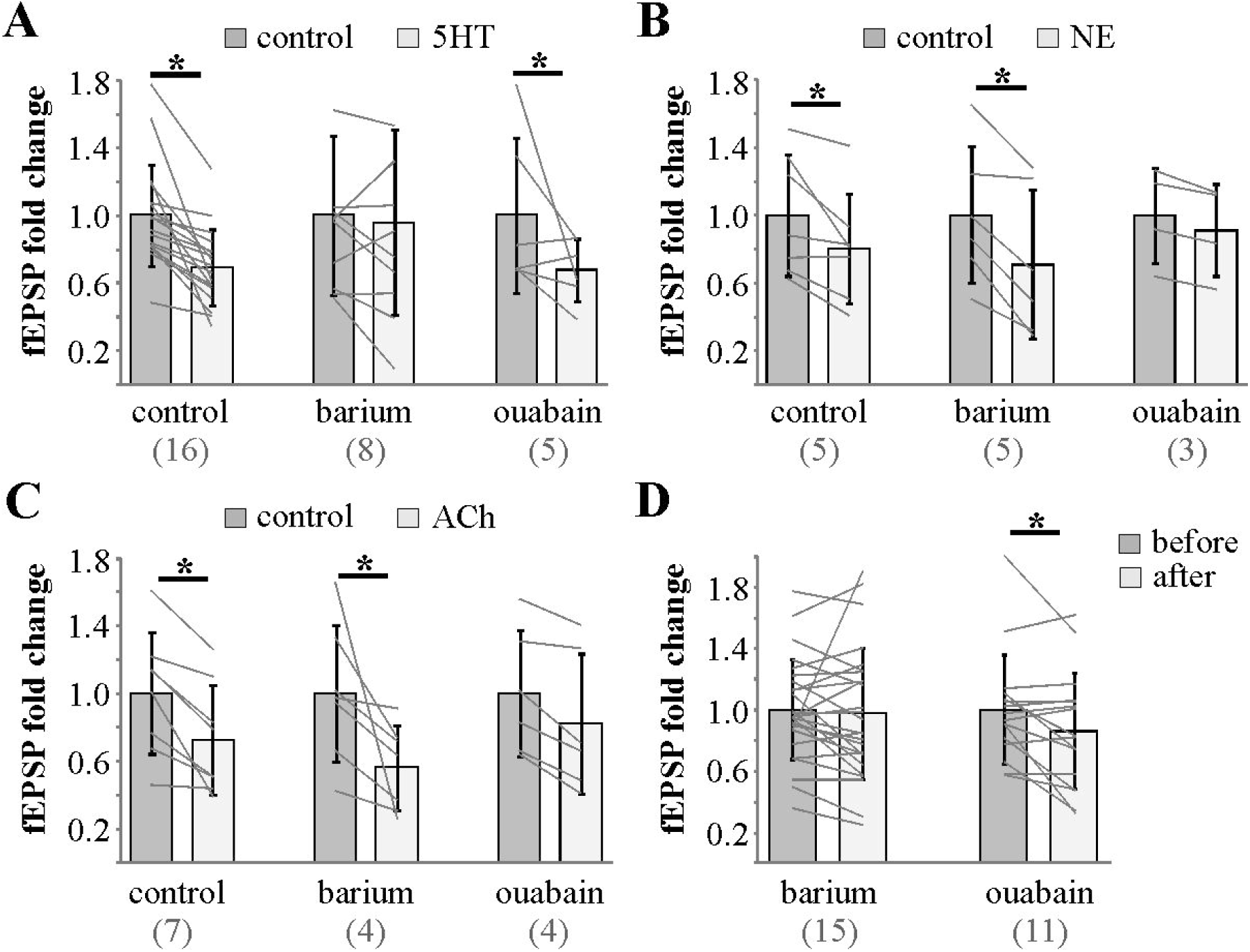
Neuromodulator-mediated effects on fEPSP amplitude are differentially affected by K^+^-directed pharmacology. **A)** The 5HT-mediated decrease in the amplitude is lost in the presence of Ba^2+^, but not ouabain. **B** and **C)** In contrast, the NE- and ACh-mediated decreases are lost in the presence of ouabain and not Ba^2+^. **D)** Only ouabain affected fEPSP amplitude upon wash-in. * p < 0.05 two-way RM ANOVA with Fisher’s LSD. n = 3-18. Numbers in parentheses on x-axis indicate the number of animals.

### Alteration of [K^+^]_e_ can directly affect synaptic adaptation and amplitude

Given that neuromodulator effects on adaptation can be disrupted with pharmacology that disrupts neuromodulator-mediated effects on [K^+^]_e_ homeostasis, we next sought to assess the impact of directly changing slice perfusate [K^+^] on adaptation in isolation of the many other likely effects neuromodulators may have on cell function. Interestingly, we only observed an effect on adaptation when we decreased perfusate [K^+^] from 3 mM to 1 mM as opposed to increasing from 3mM to 6 mM (Fig. 7A, B; potassium x time interaction p = 0.011; F_1, 16_ = 8.227). This was consistent with effects on evoked K^+^ recovery tau (Fig. 7C; potassium x time interaction p = 0.020; F_1, 10_ = 7.642). Changing the bath perfusate from 3 mM to 1 mM K^+^ also lead to a decrease in fEPSP amplitude (−0.64 ± 0.038 vs −0.43 ± 0.040 mV for 3 mM and 1 mM K^+^, respectively; P < 0.001). No effect was observed when changing to 6 mM (−0.90 ± 0.14 vs - 0.88 ± 0.16 for 3 mM and 6 mM, respectively; P = 0.69).

**Figure 7:**
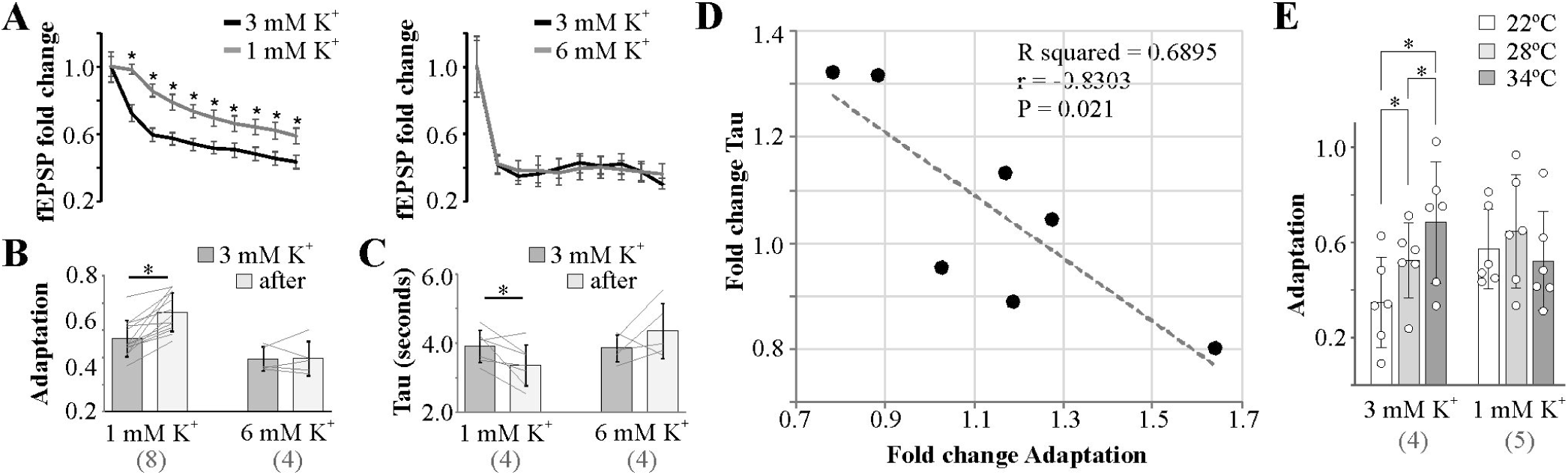
Directly decreasing K^+^ in the perfusate replicates 5HT and NE effects on adaptation and evoked K^+^ recovery tau. **A** and **B)** Reducing bath [K^+^] from 3 mM to 1 mM reduces adaptation whereas increasing from 3 mM to 6 mM has no effect. **C)** Effects on adaptation with changes in perfusate [K^+^] are reflected in evoked K^+^ recovery tau. **D)** Negative correlation observed between changes in adaptation with changes in recovery tau. **E)** The adaptation decrease observed with increasing temperature is lost when bath [K^+^] is 1 mM. * p < 0.05 two-way repeated measures ANOVA with Fisher’s LSD. n = 5-13. Numbers in parentheses on x-axis indicate the number of animals.

It is worth noting that although 5HT and NE affect baseline [K^+^] differently (Fig. 2 C-E), they both decrease evoked recovery tau and adaptation similarly (synaptic K+ accumulation), consistent with effects of lowering perfusate [K^+^] that also decreases evoked recovery tau and adaptation. In conjunction with the fact that ACh does not impact evoked recovery tau or somatosensory adaptation, we looked to see if changes to recovery tau correlated with changes in adaptation. Using data where only perfusate [K^+^] was changed, ruling out other aspects of neuromodulator-mediated receptor actions, we observe a significant negative correlation (Pearson’s r = −0.83, R^2^ = 0.69, p = 0.021) between changes in adaptation and changes in recovery tau (Fig. 7D). Thus, it appears that changes in extracellular [K^+^] regulation is sufficient to affect somatosensory adaptation. It should be noted that the lack of effect of increasing perfusate [K^+^] to 6 mM on recovery tau and adaptation is consistent with the ACh-mediated lack of effect on recovery tau and adaptation. Raising [K^+^]_e_ alone (ACh or 6 mM perfusate) has no impact on somatosensory adaptation. However, lowering [K^+^]_e_ alone replicated both 5HT- and NE-mediated effects. Although 5HT does not directly lower baseline [K^+^]_e_ (Fig. 1E), by decreasing recovery tau, 5HT and NE effectively lower or reduce K^+^ accumulation in the synapse.

The approach above replicates 5HT and NE effects on adaptation by lowering bath perfusate to indirectly reduce K^+^ accumulation in the synapse. An alternate approach would be to increase K^+^ clearance at the synapse. Although these studies demonstrate a role for both Kir4.1 and NKA activity in neuromodulator effects, we likely underestimate the role of the NKA enzyme activity as it is well known to be temperature-sensitive with NKA activity increasing at physiological temperatures (Ransom, Ransom, & Sontheimer, 2000). Thus, additional experiments were performed assessing adaptation at 22°C (room temperature), 28°C, and 34°C for both normal and low bath aCSF [K^+^] (1 mM; Fig. 7E). The rationale being that 1 mM K^+^ aCSF should occlude effects of temperature on NKA activity if the effects on adaptation are mediated via synaptic K^+^ accumulation. Indeed, we found that adaptation was reduced when temperature was increased and this temperature-mediated effect was lost in 1mM K^+^ aCSF (Fig. 7E; repeated measures two-way ANOVA temperature x potassium interaction p = 0.003; F_2, 20_ = 7.708). Thus, we show that reducing K^+^ accumulation in the synapse by 1) reducing bath perfusate K^+^ or 2) increasing NKA activity with increasing temperatures results in similar effects on adaptation, strongly supporting a direct effect of K^+^ clearance in somatosensory spike frequency adaptation.

### Disruption of glycolysis supports an astrocyte-dependent neuromodulator-mediated K^+^ regulation

Although Ba^2+^-sensitive inward rectifying K^+^ channels and ouabain-sensitive NKA activity have been consistently shown to be involved in K^+^ clearance in the hippocampus and largely a function of astrocytes (D’Ambrosio et al., 2002; Larsen et al., 2014; Larsen, Holm, et al., 2016; Macaulay & Zeuthen, 2012; Murakami & Kurachi, 2016; Sibille et al., 2015), we assessed astrocyte involvement in K^+^ clearance in the cortex using the glyceraldehyde phosphate dehydrogenase (GAPDH) inhibitor iodoacetate (IDA) to disrupt the energy producing step in glycolysis (Schmidt & Dringen, 2009)(Fig. 8A) thought to be involved in fueling astrocyte NKA activity (DiNuzzo, Mangia, Maraviglia, & Giove, 2013; Mangia, Giove, & Dinuzzo, 2013). As astrocytes rely more heavily on glycolysis (DiNuzzo, Mangia, Maraviglia, & Giove, 2010; Pellerin et al., 2007) and neurons more so on oxidative phosphorylation (Wyss, Jolivet, Buck, Magistretti, & Weber, 2011), we add lactate (1 mM) to the aCSF to help maintain neuronal metabolic needs. As such, the low-dose IDA we use results in a relatively selective disruption of astrocyte metabolism. Consistent with 5HT-mediated effects primarily involving Ba^2+^-sensitive inward rectifying K^+^ channels at room temperature, IDA (100 µM; IC_50_ = 100 µM (Schmidt & Dringen, 2009)) showed a minor effect on recovery tau (loss of decrease; Fig. 8B, top) and no effect on adaptation or fEPSP amplitude (Fig. 8B, middle, bottom). In contrast, IDA prevented NE-mediated effects on recovery tau (Fig. 8C, top; pharm x time interaction p = 0.002; F_1, 14_ = 15.19) and adaptation (Fig. 8C, middle; pharm x time interaction p = 0.004; F_1, 11_ = 13.00) with, again, no impact on fEPSP amplitude (Fig. 8C, bottom). Together, data obtained with IDA is identical to that we observed with low-dose ouabain, supporting the notion that astrocytes are at the heart of neuromodulator-mediated effects on potassium homeostasis.

**Figure 8:**
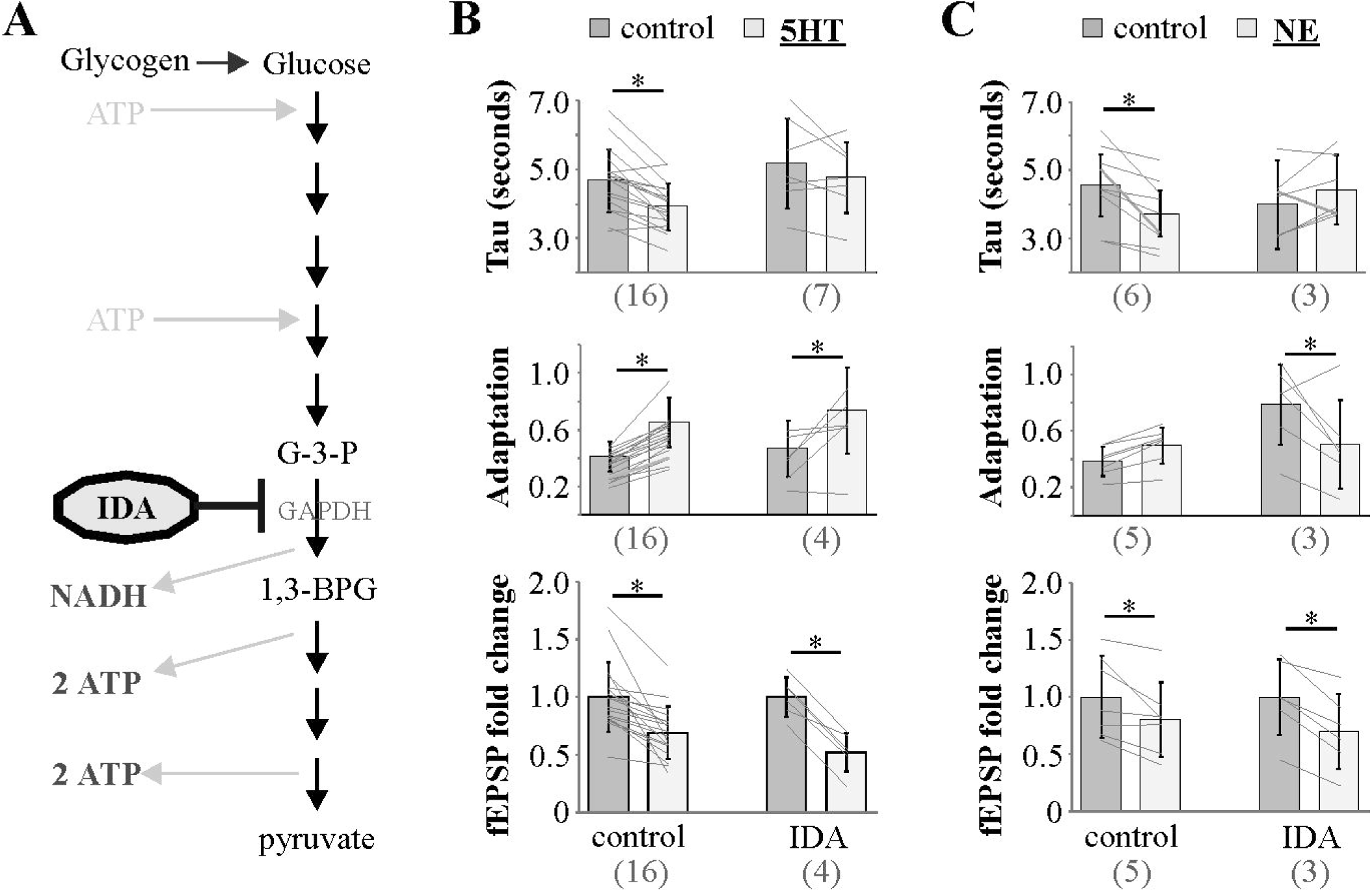
Disruption of astrocyte glycolysis supports a role for astrocytes in 5HT and NE effects. **A)** Iodoacetate (IDA) inhibits GAPDH and thereby the energy-producing reactions in glycolysis/glycogenolysis. **B)** IDA disrupts 5HT-mediated effects on recovery tau (top) but not adaptation (middle) or fEPSP amplitude (bottom). **C)** IDA disrupts NE-mediated effects on recovery tau (top) and adaptation (middle), but not fEPSP amplitude (bottom). * p < 0.05 two-way repeated measures ANOVA with Fisher’s LSD. n = 6-16. Numbers in parentheses on x-axis indicate the number of animals.

## Discussion

This is the first study to implicate astrocyte-mediated regulation of extracellular potassium in neuromodulator-mediated effects on sensory transmission in the cortex. Given that individual astrocytes ensheath up to 140,000 synapses (Bushong, Martone, & Ellisman, 2004; Bushong, Martone, Jones, & Ellisman, 2002) (both excitatory and inhibitory) and each regulate the environment of their own non-overlapping territory (Oberheim, Goldman, & Nedergaard, 2012; Oberheim et al., 2008; Oberheim, Wang, Goldman, & Nedergaard, 2006), astrocyte-mediated regulation of [K^+^]_e_ would serve a highly efficient means by which diffuse, long-range neuromodulatory networks (5HT, NE and ACh from raphe, locus coeruleus and nucleus basalis of Meynert, respectively) could rapidly affect broad sensory fields. Consistent with this notion, we demonstrate that 5HT, NE and ACh differentially affect [K^+^]_e_ regulation (Fig. 2) via the Kir4.1 or NKA (Fig. 3) that mirrors differential neuromodulatory effects on somatosensory adaptation (Figs. 4/5) and amplitude (Fig. 6) (summarized in Fig. 9).

**Figure 9:**
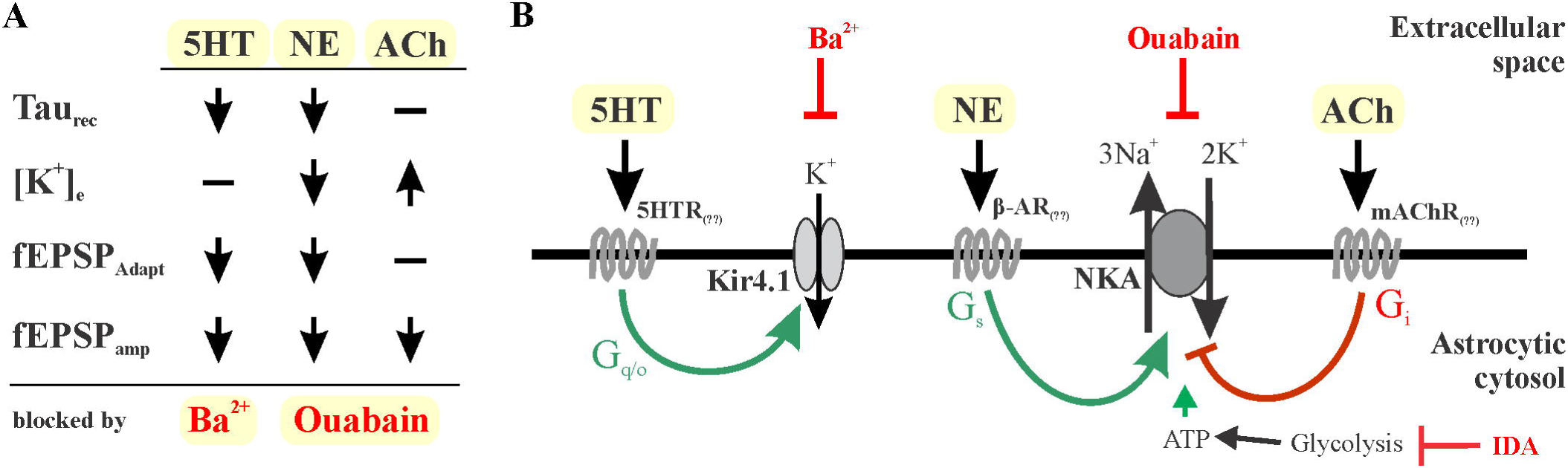
Neuromodulator-mediated effects are sensitive to block of Kir4.1 or NKA. **A)** Neuromodulator-specific effects on extracellular potassium management, spike frequency adaptation and fEPSP amplitude. **B)** All 5HT effects measured in this study are sensitive to Ba^2+^-mediated block of inward rectifying potassium channels. All NE- and ACh-mediated effects are sensitive to ouabain-mediated suppression of NKA activity. Ouabain blocks stimulation with respect to NE and blocks suppression with respect to ACh.

Clearance of synaptic K^+^ has consistently been attributed to astrocytes (D’Ambrosio et al., 2002; Kofuji & Newman, 2004; Larsen et al., 2014; Larsen, Holm, et al., 2016; Larsen, Stoica, et al., 2016; Macaulay & Zeuthen, 2012; Murakami & Kurachi, 2016; Ransom et al., 2000; Sibille et al., 2015). The α2β2 subunit of the NKA, specific to astrocytes (Cameron, Klein, Shyjan, Rakic, & Levenson, 1994; Larsen, Stoica, et al., 2016; Zhang et al., 2014; Zhang et al., 2016), is the only subunit not saturated under physiological K^+^ concentrations and therefore demonstrates activity increases in response to synaptic K^+^ transients (Larsen et al., 2014). α1 and α3 subunits in neurons contribute at rest, but activity does not change with synaptic activity and thus astrocytes are responsible for the early evoked K^+^ clearance (Larsen, Holm, et al., 2016; Larsen, Stoica, et al., 2016). Inwardly rectifying K^+^ channels are also important for spatial buffering of K^+^ across voltage gradients within the astrocyte syncytium (Larsen & MacAulay, 2014; Murakami & Kurachi, 2016). Neurons have inward rectifying K^+^ channels, but membrane potentials are too depolarized to develop inward K^+^ fluxes. The more negative membrane potential and isopotentiality (Kiyoshi et al., 2018; Kiyoshi & Zhou, 2019) of the astrocyte syncytium creates the ideal scenario for inward rectifier-mediated buffering of local synaptic K^+^ transients. Disruption of glycolysis with iodoacetate in these studies provide additional support for astrocyte involvement in neuromodulator-mediated effects. Disruption of glycolysis in the presence of lactate has minimal effect on neurons. However, astrocytes, being the sole source of glycogen in the brain, rely heavily on glycolysis for energy to fuel the NKA (DiNuzzo et al., 2013; Mangia et al., 2013). In this regard, it is not surprising that IDA experiments replicated the results obtained using the NKA inhibitor ouabain.

The different neuromodulators can indirectly affect Kir4.1 or NKA via 2^nd^ messenger signaling cascades. Regulation of NKA activity appears to be prominently dependent on cAMP levels that affect protein kinase A-mediated phosphorylation of the NKA α-subunit(Bertorello & Katz, 1993; Ewart & Klip, 1995). Studies in astrocyte cultures have demonstrated that NE stimulates NKA activity via β-adrenergic and adenylyl cyclase activation(Hajek, Subbarao, & Hertz, 1996; Hertz & Chen, 2016). Protein kinase C and calmodulin-kinase can also affect NKA activity and/or expression at the cell surface(Bertorello & Katz, 1993; Ewart & Klip, 1995; Gao et al., 1999). Acetylcholine muscarinic receptors are known to inhibit adenylyl cyclase and reduce cAMP levels to inhibit β-agonist effects on NKA activity(Gao, Mathias, Cohen, & Baldo, 1997). Another interesting aspect of NKA regulation is the dependence on intracellular [Na^+^](Efendiev, Bertorello, Zandomeni, Cinelli, & Pedemonte, 2002) as it is not high enough at rest to saturate enzyme activity. Synaptic activity can drive Na^+^ increase in astrocytes via the glutamate transporter to increase NKA activity(Larsen, Holm, et al., 2016), but neuromodulators often function independently of active synapses and so need another means to provide Na^+^ for fueling NKA activity. In this regard, it is no coincidence that astrocyte calcium transients can drive intracellular Na^+^ increases via the Na^+^-Ca^2+^ exchanger(Wang, Smith, et al., 2012) and that astrocytes show robust calcium responses to the different neuromodulators(Bekar, He, & Nedergaard, 2008; Ding et al., 2013; Perea & Araque, 2005; Schipke, Heuser, & Peters, 2011; Shelton & McCarthy, 2000). Regulation of inward rectifying potassium channels in astrocytes is also mediated by protein kinases and phosphatases. Protein kinase A increases(Bolton, Greenwood, Hamilton, & Butt, 2006) and protein kinase C decreases(Koller, Allert, Oel, Stoll, & Siebler, 1998) inward rectifier conductance in astrocytes. 5HT has been shown to increase inward rectifying current in CA3 neurons via 5HT_1A_ receptors(Okuhara & Beck, 1994) as well as decrease potassium conductance in nucleus accumbens neurons via 5HT_2_ receptors(North & Uchimura, 1989). Thus, the ultimate effect on Kir4.1 or NKA activity is determined by the receptor complement that may be cell and/or region specific.

Large experimentally evoked K^+^ responses typically show a rapid increase followed by a decrease/recovery phase that often demonstrates an undershoot before returning to baseline [K^+^]. Although it is commonly accepted that both Kir4.1 and NKA activity are involved in regulation of evoked K^+^ responses (D’Ambrosio et al., 2002; Larsen et al., 2014; Sibille et al., 2014), the role of Kir4.1 channels have only figured into the evoked amplitude (Larsen et al., 2014) or recovery undershoot (Chever et al., 2010; D’Ambrosio et al., 2002) with no impact on rate of recovery (D’Ambrosio et al., 2002; Larsen et al., 2014; Sibille et al., 2014). The rate of recovery has been attributed most prominently to NKA activity that can be driven by glutamate transporter-mediated intracellular Na^+^ transients (Hertz & Chen, 2016; Larsen, Holm, et al., 2016; Sibille et al., 2014). However, as it is suggested that Kir4.1-mediated uptake requires non-uniform distribution of K^+^ increases (Larsen & MacAulay, 2014; Murakami & Kurachi, 2016; Sibille et al., 2015), large/global K^+^ increases could limit the observation of Kir4.1 in the spatially-restricted recording of evoked K^+^ recovery. Previous studies assessing [K^+^]_e_ typically used long-lasting stimulation protocols (200-900 stimuli; 1-20 Hz)(D’Ambrosio et al., 2002; Larsen et al., 2014) that lead to large, almost global, increases in K^+^ that limit the gradient necessary to observe Kir4.1 involvement in K^+^ clearance/removal (Larsen & MacAulay, 2014). We propose that the short-lasting stimulation (10 stimuli; 20 Hz) used in this study that elicited a small local K^+^ increase (∼0.5 mM), maintained gradients for observation of changes in evoked recovery in response to 5HT and NE that were sensitive to Ba^2+^ blockade. Similar short-lasting stimulation (5-20 stimuli; 10 Hz) in normal and Kir4.1 cKO mouse hippocampus *in vivo* demonstrated significant dependence on Kir4.1 channels (Chever et al., 2010), supporting the concept that under typical firing patterns, spatial buffering through Kir4.1 is instrumental in maintaining synaptic [K^+^]_e_.

The sensitivity of neuromodulator-mediated recovery changes (5HT and NE) to both Ba^2+^ and ouabain in this study suggest the two mechanisms may be interdependent in establishing a gradient for spatial buffering. We had expected the NKA to be the main mediator of the increase in recovery (D’Ambrosio et al., 2002; Hertz & Chen, 2016; Larsen et al., 2014), so were surprised to find that Ba^2+^ had a similar inhibitory effect on both the 5HT- and NE-mediated increase in recovery. It may be that both perfusate applied Ba^2+^ and ouabain can globally depolarize astrocytes (Chever et al., 2010; Meeks & Mennerick, 2007), thereby disrupting the gradient for K^+^ spatial buffering across the glial syncytium. Combined with the limited spatial resolution of extracellular K^+^ ion-selective recording, subtle changes at the synaptic level are no longer detectable. However, effects on synaptic adaptation observed in this study (averaged effect at the spatially-restricted synaptic level) supported the differential (Ba^2+^-versus ouabain-sensitive) neuromodulator effect on local spatially-restricted synaptic K^+^ uptake. Thus, lack of Ba^2+^/ouabain selectivity for the different neuromodulators in the non-spatially-restricted ion-selective recordings reflects a potential dissolution of global K^+^ gradients necessary for evoked K^+^ recovery. On the other hand, potassium uptake at the spatially-restricted synaptic level may explain the observed selectivity maintained in our synaptic adaptation studies. The fact that neither Ba^2+^ nor 5HT affect cortical baseline potassium levels in the presence of TTX, supports the notion that the 5HT-mediated increase in evoked K^+^ recovery does not involve the NKA despite being disrupted by ouabain (ouabain did affect baseline). Both NE and ACh effects on baseline [K^+^]_e_, however, did involve effects on NKA activity as they were inhibited or occluded by low-dose ouabain, respectively. The fact that ACh- or ouabain-mediated increase in [K^+^]_e_ does not affect evoked K^+^ recovery suggests NKA activity in these studies is already very low with the mild stimulation used. Such a scenario would account for why NE-mediated stimulation of NKA affects evoked K^+^ recovery whereas ACh-mediated inhibition has no effect (already negligible).

Both NE and 5HT decreased evoked fEPSP amplitude and frequency adaptation whereas acetylcholine only decreased the amplitude. Although NE and 5HT effects appear similar under conditions used in our studies, the NE effects were abolished by the NKA inhibitor ouabain while 5HT effects were abolished by the Kir inhibitor Ba^2+^ (Fig. 9A). The lack of obvious differences likely relates to the output measures used in this study. Compared to extracellular field recordings, patch-clamp analysis of a single post-synaptic neuron may enable improved ability to differentiate the neuromodulator responses. Nonetheless, the pharmacology of the effects is clearly different. Disruption of ACh-mediated effects on [K^+^]_e_ with ouabain also disrupted effects on fEPSP amplitude supporting the notion that [K^+^]_e_, not receptor transduction mechanisms, is the effector. Consistent with this, decreasing perfusate K^+^ from 3 mM to 1 mM was able to recapitulate NE and 5HT effects on adaptation as well as amplitude. Thus, decreasing spatially restricted synaptic K^+^, no matter the mechanism, via NKA, Kir, or 1mM perfusate, all results in similar effects on fEPSP frequency adaptation and amplitude.

Evidence now exists that supports a role for regulation of potassium homeostasis in both healthy and pathological brain neuromodulation. Extracellular potassium can reach concentrations of 10 - 12 mM with seizures and as high as 80 mM during ischemia or spreading depression (Hertz & Chen, 2016); conditions that help suppress neural activity. Under physiological conditions, maintenance of extracellular potassium has been implicated in long-term plasticity (D’Ambrosio, Maris, Grady, Winn, & Janigro, 1998; Janigro, Gasparini, D’Ambrosio, McKhann, & DiFrancesco, 1997) as well as sleep and behavior (Ding et al., 2016). Calcium-dependent K^+^ uptake in glial cells has also been shown to affect the signal-to-noise ratio of synaptic transmission in the hippocampus (Wang, Smith, et al., 2012) and modulate bistability (up/down-state) of Purkinje cells in the cerebellum (Wang, Xu, Wang, Takano, & Nedergaard, 2012). Calcium events in astrocytes have also been shown to regulate cortical bistability (Poskanzer & Yuste, 2011, 2016). Considering the impact that [K^+^]_e_ can have on many neural and circuit functions; it seems entirely reasonable that neuromodulators regulate extracellular potassium to exert their many effects. This study provides functional significance for astrocyte-mediated buffering of extracellular potassium in neuromodulator-mediated shaping of cortical network activity.

## Supporting information

Manuscript data file

## Conflict of Interest Statement

The authors declare no conflicts of interest.

## Author Contributions

All authors had full access to all the data in the study and take responsibility for the integrity of the data and the accuracy of the data analysis.

*Conceptualization*, L.K.B. and C.A.W.; *Methodology*, L.K.B. and C.A.W.; *Investigation*, C.A.W. and C.D.C.; *Formal Analysis*, C.A.W. and C.D.C.; *Writing – Original Draft*, L.K.B. and C.A.W.; *Writing – Review & Editing*, L.K.B., C.A.W. and C.D.C.; *Visualization*, L.K.B. and C.A.W.; *Supervision*, L.K.B.; *Funding Acquisition*, L.K.B.

## Acknowledgments

This work was supported by the Natural Sciences and Engineering Research Council of Canada (NSERC; grant number 195814317).

## Data availability Statement

The data that support the findings of this study are available as a supplementary data file (Microsoft excel).

## Notes

**Grant Support:** This work was supported by the Natural Sciences and Engineering Research Council of Canada (NSERC; grant number 195814317).

## References

Abbott, L. F., Varela, J. A., Sen, K., & Nelson, S. B. (1997). Synaptic depression and cortical gain control. Science, 275(5297), 220–224.

Ammann, D., Chao, P. S., & Simon, W. (1987). Valinomycin-based K+ selective microelectrodes with low electrical membrane resistance. Neurosci Lett, 74(2), 221–226. doi:10.1016/0304-3940(87)90153-4

Bekar, L. K., He, W., & Nedergaard, M. (2008). Locus coeruleus alpha-adrenergic-mediated activation of cortical astrocytes in vivo. Cerebral cortex, 18(12), 2789–2795. doi:10.1093/cercor/bhn040

Bellot-Saez, A., Kekesi, O., Morley, J. W., & Buskila, Y. (2017). Astrocytic modulation of neuronal excitability through K(+) spatial buffering. Neurosci Biobehav Rev, 77, 87–97. doi:10.1016/j.neubiorev.2017.03.002

Bertorello, A. M., & Katz, A. I. (1993). Short-term regulation of renal Na-K-ATPase activity: physiological relevance and cellular mechanisms. Am J Physiol, 265(6 Pt 2), F743–755. doi:10.1152/ajprenal.1993.265.6.F743

Bolton, S., Greenwood, K., Hamilton, N., & Butt, A. M. (2006). Regulation of the astrocyte resting membrane potential by cyclic AMP and protein kinase A. Glia, 54(4), 316–328. doi:10.1002/glia.20384

Bushong, E. A., Martone, M. E., & Ellisman, M. H. (2004). Maturation of astrocyte morphology and the establishment of astrocyte domains during postnatal hippocampal development. Int J Dev Neurosci, 22(2), 73–86. doi:10.1016/j.ijdevneu.2003.12.008

Bushong, E. A., Martone, M. E., Jones, Y. Z., & Ellisman, M. H. (2002). Protoplasmic astrocytes in CA1 stratum radiatum occupy separate anatomical domains. J Neurosci, 22(1), 183–192.

Butt, A. M., & Kalsi, A. (2006). Inwardly rectifying potassium channels (Kir) in central nervous system glia: a special role for Kir4.1 in glial functions. J Cell Mol Med, 10(1), 33–44. doi:10.1111/j.1582-4934.2006.tb00289.x

Cameron, R., Klein, L., Shyjan, A. W., Rakic, P., & Levenson, R. (1994). Neurons and astroglia express distinct subsets of Na,K-ATPase alpha and beta subunits. Brain Res Mol Brain Res, 21(3-4), 333–343. doi:10.1016/0169-328x(94)90264-x

Castro-Alamancos, M. A., & Gulati, T. (2014). Neuromodulators produce distinct activated states in neocortex. J Neurosci, 34(37), 12353–12367. doi:10.1523/jneurosci.1858-14.2014

Chever, O., Djukic, B., McCarthy, K. D., & Amzica, F. (2010). Implication of Kir4.1 channel in excess potassium clearance: an in vivo study on anesthetized glial-conditional Kir4.1 knock-out mice. J Neurosci, 30(47), 15769–15777. doi: 30/47/15769 [pii] 10.1523/JNEUROSCI.2078-10.2010

D’Ambrosio, R., Gordon, D. S., & Winn, H. R. (2002). Differential role of KIR channel and Na(+)/K(+)-pump in the regulation of extracellular K(+) in rat hippocampus. J Neurophysiol, 87(1), 87–102.

D’Ambrosio, R., Maris, D. O., Grady, M. S., Winn, H. R., & Janigro, D. (1998). Selective loss of hippocampal long-term potentiation, but not depression, following fluid percussion injury. Brain Res, 786(1-2), 64–79.

Deng, P. Y., & Lei, S. (2008). Serotonin increases GABA release in rat entorhinal cortex by inhibiting interneuron TASK-3 K+ channels. Mol Cell Neurosci, 39(2), 273–284. doi:10.1016/j.mcn.2008.07.005

Ding, F., O’Donnell, J., Thrane, A. S., Zeppenfeld, D., Kang, H., Xie, L., … Nedergaard, M. (2013). alpha1-Adrenergic receptors mediate coordinated Ca2+ signaling of cortical astrocytes in awake, behaving mice. Cell Calcium, 54(6), 387–394. doi:10.1016/j.ceca.2013.09.001

Ding, F., O’Donnell, J., Xu, Q., Kang, N., Goldman, N., & Nedergaard, M. (2016). Changes in the composition of brain interstitial ions control the sleep-wake cycle. Science, 352(6285), 550–555. doi:10.1126/science.aad4821

DiNuzzo, M., Mangia, S., Maraviglia, B., & Giove, F. (2010). Glycogenolysis in astrocytes supports blood-borne glucose channeling not glycogen-derived lactate shuttling to neurons: evidence from mathematical modeling. J Cereb Blood Flow Metab, 30(12), 1895–1904. doi:10.1038/jcbfm.2010.151

DiNuzzo, M., Mangia, S., Maraviglia, B., & Giove, F. (2013). Regulatory mechanisms for glycogenolysis and K+ uptake in brain astrocytes. Neurochem Int, 63(5), 458–464. doi:10.1016/j.neuint.2013.08.004

Efendiev, R., Bertorello, A. M., Zandomeni, R., Cinelli, A. R., & Pedemonte, C. H. (2002). Agonist-dependent regulation of renal Na+,K+-ATPase activity is modulated by intracellular sodium concentration. J Biol Chem, 277(13), 11489–11496. doi:10.1074/jbc.M108182200

Ewart, H. S., & Klip, A. (1995). Hormonal regulation of the Na(+)-K(+)-ATPase: mechanisms underlying rapid and sustained changes in pump activity. Am J Physiol, 269(2 Pt 1), C295–311.

Fioravante, D., & Regehr, W. G. (2011). Short-term forms of presynaptic plasticity. Curr Opin Neurobiol, 21(2), 269–274. doi:10.1016/j.conb.2011.02.003

Gao, J., Mathias, R. T., Cohen, I. S., & Baldo, G. J. (1997). Effects of acetylcholine on the Na(+)-K+ pump current in guinea-pig ventricular myocytes. J Physiol, 501 (Pt 3), 527–535. doi:10.1111/j.1469-7793.1997.527bm.x

Gao, J., Mathias, R. T., Cohen, I. S., Wang, Y., Sun, X., & Baldo, G. J. (1999). Activation of PKC increases Na+-K+ pump current in ventricular myocytes from guinea pig heart. Pflugers Arch, 437(5), 643–651. doi:10.1007/s004240050828

Hajek, I., Subbarao, K. V., & Hertz, L. (1996). Acute and chronic effects of potassium and noradrenaline on Na+, K+-ATPase activity in cultured mouse neurons and astrocytes. Neurochem Int, 28(3), 335–342.

Hertz, L., & Chen, Y. (2016). Importance of astrocytes for potassium ion (K(+)) homeostasis in brain and glial effects of K(+) and its transporters on learning. Neurosci Biobehav Rev, 71, 484–505. doi:10.1016/j.neubiorev.2016.09.018

Hirata, A., Aguilar, J., & Castro-Alamancos, M. A. (2006). Noradrenergic activation amplifies bottom-up and top-down signal-to-noise ratios in sensory thalamus. J Neurosci, 26(16), 4426–4436.

Janigro, D., Gasparini, S., D’Ambrosio, R., McKhann, G., 2nd, & DiFrancesco, D. (1997). Reduction of K+ uptake in glia prevents long-term depression maintenance and causes epileptiform activity. J Neurosci, 17(8), 2813–2824.

Kiyoshi, C. M., Du, Y., Zhong, S., Wang, W., Taylor, A. T., Xiong, B., … Zhou, M. (2018). Syncytial isopotentiality: A system-wide electrical feature of astrocytic networks in the brain. Glia, 66(12), 2756–2769. doi:10.1002/glia.23525

Kiyoshi, C. M., & Zhou, M. (2019). Astrocyte syncytium: a functional reticular system in the brain. Neural Regen Res, 14(4), 595–596. doi:10.4103/1673-5374.247462

Kofuji, P., & Newman, E. A. (2004). Potassium buffering in the central nervous system. Neuroscience, 129(4), 1045–1056. doi:10.1016/j.neuroscience.2004.06.008

Koller, H., Allert, N., Oel, D., Stoll, G., & Siebler, M. (1998). TNF alpha induces a protein kinase C-dependent reduction in astroglial K+ conductance. Neuroreport, 9(7), 1375–1378. doi:10.1097/00001756-199805110-00023

Larsen, B. R., Assentoft, M., Cotrina, M. L., Hua, S. Z., Nedergaard, M., Kaila, K., … MacAulay, N. (2014). Contributions of the Na(+)/K(+)-ATPase, NKCC1, and Kir4.1 to hippocampal K(+) clearance and volume responses. Glia, 62(4), 608–622. doi:10.1002/glia.22629

Larsen, B. R., Holm, R., Vilsen, B., & MacAulay, N. (2016). Glutamate transporter activity promotes enhanced Na(+) /K(+) -ATPase-mediated extracellular K(+) management during neuronal activity. J Physiol, 594(22), 6627–6641. doi:10.1113/jp272531

Larsen, B. R., & MacAulay, N. (2014). Kir4.1-mediated spatial buffering of K(+): experimental challenges in determination of its temporal and quantitative contribution to K(+) clearance in the brain. Channels (Austin), 8(6), 544–550. doi:10.4161/19336950.2014.970448

Larsen, B. R., Stoica, A., & MacAulay, N. (2016). Managing Brain Extracellular K(+) during Neuronal Activity: The Physiological Role of the Na(+)/K(+)-ATPase Subunit Isoforms. Front Physiol, 7, 141. doi:10.3389/fphys.2016.00141

Lei, S., Deng, P. Y., Porter, J. E., & Shin, H. S. (2007). Adrenergic facilitation of GABAergic transmission in rat entorhinal cortex. J Neurophysiol, 98(5), 2868–2877. doi: 00679.2007 [pii] 10.1152/jn.00679.2007

Macaulay, N., & Zeuthen, T. (2012). Glial K(+) clearance and cell swelling: key roles for cotransporters and pumps. Neurochem Res, 37(11), 2299–2309. doi:10.1007/s11064-012-0731-3

Mangia, S., Giove, F., & Dinuzzo, M. (2013). K+ homeostasis in the brain: a new role for glycogenolysis. Neurochem Res, 38(3), 470–471. doi:10.1007/s11064-012-0962-3

McCormick, D. A. (1992). Neurotransmitter actions in the thalamus and cerebral cortex and their role in neuromodulation of thalamocortical activity. Prog Neurobiol, 39(4), 337–388. doi:0301-0082(92)90012-4 [pii]

Meeks, J. P., & Mennerick, S. (2007). Astrocyte membrane responses and potassium accumulation during neuronal activity. Hippocampus, 17(11), 1100–1108. doi:10.1002/hipo.20344

Murakami, S., & Kurachi, Y. (2016). Mechanisms of astrocytic K(+) clearance and swelling under high extracellular K(+) concentrations. J Physiol Sci, 66(2), 127–142. doi:10.1007/s12576-015-0404-5

North, R. A., & Uchimura, N. (1989). 5-Hydroxytryptamine acts at 5-HT2 receptors to decrease potassium conductance in rat nucleus accumbens neurones. J Physiol, 417, 1–12. doi:10.1113/jphysiol.1989.sp017786

Oberheim, N. A., Goldman, S. A., & Nedergaard, M. (2012). Heterogeneity of astrocytic form and function. Methods Mol Biol, 814, 23–45. doi:10.1007/978-1-61779-452-0_3

Oberheim, N. A., Tian, G. F., Han, X., Peng, W., Takano, T., Ransom, B., & Nedergaard, M. (2008). Loss of astrocytic domain organization in the epileptic brain. J Neurosci, 28(13), 3264–3276. doi:10.1523/jneurosci.4980-07.2008

Oberheim, N. A., Wang, X., Goldman, S., & Nedergaard, M. (2006). Astrocytic complexity distinguishes the human brain. Trends Neurosci, 29(10), 547–553.

Okuhara, D. Y., & Beck, S. G. (1994). 5-HT1A receptor linked to inward-rectifying potassium current in hippocampal CA3 pyramidal cells. J Neurophysiol, 71(6), 2161–2167. doi:10.1152/jn.1994.71.6.2161

Pellerin, L., Bouzier-Sore, A. K., Aubert, A., Serres, S., Merle, M., Costalat, R., & Magistretti, P. J. (2007). Activity-dependent regulation of energy metabolism by astrocytes: an update. Glia, 55(12), 1251–1262. doi:10.1002/glia.20528

Pellerin, L., & Magistretti, P. J. (1997). Glutamate uptake stimulates Na+,K+-ATPase activity in astrocytes via activation of a distinct subunit highly sensitive to ouabain. J Neurochem, 69(5), 2132–2137. doi:10.1046/j.1471-4159.1997.69052132.x

Perea, G., & Araque, A. (2005). Properties of synaptically evoked astrocyte calcium signal reveal synaptic information processing by astrocytes. J Neurosci, 25(9), 2192–2203.

Poskanzer, K. E., & Yuste, R. (2011). Astrocytic regulation of cortical UP states. Proc Natl Acad Sci U S A, 108(45), 18453–18458. doi:10.1073/pnas.1112378108

Poskanzer, K. E., & Yuste, R. (2016). Astrocytes regulate cortical state switching in vivo. Proc Natl Acad Sci U S A, 113(19), E2675–2684. doi:10.1073/pnas.1520759113

Quon, E. F., Wotton, C. A., & Bekar, L. K. (2017). Evidence for astrocyte purinergic signaling in cortical sensory adaptation and serotonin-mediated neuromodulation. Mol Cell Neurosci, 88, 53–61. doi:10.1016/j.mcn.2017.12.008

Ransom, C. B., Ransom, B. R., & Sontheimer, H. (2000). Activity-dependent extracellular K+ accumulation in rat optic nerve: the role of glial and axonal Na+ pumps. J Physiol, 522 Pt 3, 427–442. doi:10.1111/j.1469-7793.2000.00427.x

Rasmussen, R., Nicholas, E., Petersen, N. C., Dietz, A. G., Xu, Q., Sun, Q., & Nedergaard, M. (2019). Cortex-wide Changes in Extracellular Potassium Ions Parallel Brain State Transitions in Awake Behaving Mice. Cell Rep, 28(5), 1182-1194.e1184. doi:10.1016/j.celrep.2019.06.082

Salgado, H., Garcia-Oscos, F., Patel, A., Martinolich, L., Nichols, J. A., Dinh, L., … Atzori, M. (2011). Layer-specific noradrenergic modulation of inhibition in cortical layer II/III. Cereb Cortex, 21(1), 212–221. doi:10.1093/cercor/bhq081

Schipke, C. G., Heuser, I., & Peters, O. (2011). Antidepressants act on glial cells: SSRIs and serotonin elicit astrocyte calcium signaling in the mouse prefrontal cortex. J Psychiatr Res, 45(2), 242–248. doi:10.1016/j.jpsychires.2010.06.005

Schmidt, M. M., & Dringen, R. (2009). Differential effects of iodoacetamide and iodoacetate on glycolysis and glutathione metabolism of cultured astrocytes. Front Neuroenergetics, 1, 1. doi:10.3389/neuro.14.001.2009

Shelton, M. K., & McCarthy, K. D. (2000). Hippocampal astrocytes exhibit Ca2+-elevating muscarinic cholinergic and histaminergic receptors in situ. J Neurochem, 74(2), 555–563.

Sibille, J., Dao Duc, K., Holcman, D., & Rouach, N. (2015). The neuroglial potassium cycle during neurotransmission: role of Kir4.1 channels. PLoS Comput Biol, 11(3), e1004137. doi:10.1371/journal.pcbi.1004137

Sibille, J., Pannasch, U., & Rouach, N. (2014). Astroglial potassium clearance contributes to short-term plasticity of synaptically evoked currents at the tripartite synapse. J Physiol, 592(1), 87–102. doi:10.1113/jphysiol.2013.261735

Stoica, A., Larsen, B. R., Assentoft, M., Holm, R., Holt, L. M., Vilhardt, F., … MacAulay, N. (2017). The alpha2beta2 isoform combination dominates the astrocytic Na(+) /K(+) -ATPase activity and is rendered nonfunctional by the alpha2.G301R familial hemiplegic migraine type 2-associated mutation. Glia, 65(11), 1777–1793. doi:10.1002/glia.23194

Sweadner, K. J. (1989). Isozymes of the Na+/K+-ATPase. Biochim Biophys Acta, 988(2), 185–220. doi:10.1016/0304-4157(89)90019-1

Varela, J. A., Sen, K., Gibson, J., Fost, J., Abbott, L. F., & Nelson, S. B. (1997). A quantitative description of short-term plasticity at excitatory synapses in layer 2/3 of rat primary visual cortex. J Neurosci, 17(20), 7926–7940.

Wang, F., Smith, N. A., Xu, Q., Fujita, T., Baba, A., Matsuda, T., … Nedergaard, M. (2012). Astrocytes modulate neural network activity by Ca^2^+-dependent uptake of extracellular K+. Sci Signal, 5(218), ra26. doi: 5/218/ra26 [pii]10.1126/scisignal.2002334

Wang, F., Xu, Q., Wang, W., Takano, T., & Nedergaard, M. (2012). Bergmann glia modulate cerebellar Purkinje cell bistability via Ca2+-dependent K+ uptake. Proc Natl Acad Sci U S A, 109(20), 7911–7916. doi:10.1073/pnas.1120380109

Wotton, C. A., Quon, E. F., Palmer, A. C., & Bekar, L. K. (2018). Corticosterone and serotonin similarly influence GABAergic and purinergic pathways to affect cortical inhibitory networks. J Neuroendocrinol, 30(4), e12592. doi:10.1111/jne.12592

Wyss, M. T., Jolivet, R., Buck, A., Magistretti, P. J., & Weber, B. (2011). In vivo evidence for lactate as a neuronal energy source. J Neurosci, 31(20), 7477–7485. doi: 31/20/7477 [pii]10.1523/JNEUROSCI.0415-11.2011

Xiao, Z., Deng, P. Y., Yang, C., & Lei, S. (2009). Modulation of GABAergic transmission by muscarinic receptors in the entorhinal cortex of juvenile rats. J Neurophysiol, 102(2), 659–669. doi:10.1152/jn.00226.2009

Zhang, Y., Chen, K., Sloan, S. A., Bennett, M. L., Scholze, A. R., O’Keeffe, S., … Wu, J. Q. (2014). An RNA-sequencing transcriptome and splicing database of glia, neurons, and vascular cells of the cerebral cortex. J Neurosci, 34(36), 11929–11947. doi:10.1523/jneurosci.1860-14.2014

Zhang, Y., Sloan, S. A., Clarke, L. E., Caneda, C., Plaza, C. A., Blumenthal, P. D., … Barres, B. A. (2016). Purification and Characterization of Progenitor and Mature Human Astrocytes Reveals Transcriptional and Functional Differences with Mouse. Neuron, 89(1), 37–53. doi:10.1016/j.neuron.2015.11.013

Zucker, R. S., & Regehr, W. G. (2002). Short-term synaptic plasticity. Annu Rev Physiol, 64, 355–405. doi:10.1146/annurev.physiol.64.092501.114547

